# Age-related differences in functional network segregation in the context of sex and reproductive stage

**DOI:** 10.1101/2022.03.28.486067

**Authors:** Hannah K. Ballard, T. Bryan Jackson, Abigail C. Miller, Tracey H. Hicks, Jessica A. Bernard

**Affiliations:** Texas A&M Institute for Neuroscience, Texas A&M University, College Station, TX, USA; Department of Psychological & Brain Sciences, Texas A&M University, College Station, TX, USA

**Keywords:** aging, functional connectivity, menopause, network segregation, sex differences

## Abstract

Age is accompanied by differences in the organization of functional brain networks, which impact behavior in adulthood. Functional networks tend to become less segregated and more integrated with age. However, sex differences in network segregation declines with age are not well-understood. Further, network segregation in the context of female reproductive stage is relatively understudied, though unmasking such relationships would be informative for elucidating biological mechanisms that contribute to sex-specific differences in aging. In the current work, we used data from the Cambridge Centre for Ageing and Neuroscience (Cam-CAN) repository to evaluate differences in resting-state network segregation as a product of sex and reproductive stage. Reproductive stage was categorized using the Stages of Reproductive Aging Workshop (STRAW+10) criteria. Replicating prior work, we investigated the following functional networks: auditory, cerebellar-basal ganglia, cingulo-opercular task control, default mode, dorsal attention, fronto-parietal task control, salience, sensory somatomotor mouth, sensory somatomotor hand, ventral attention, and visual. First, our results mirror findings from previous work indicating that network segregation is lower with increasing age. Second, when analyzing associations between network segregation and age within each sex separately, we find differences between females and males. Finally, we report significant effects of reproductive stage on network segregation, though these findings are likely driven by age. Broadly, our results suggest that impacts of sex are important to evaluate when investigating network segregation differences across adulthood, though further work is needed to determine the unique role of menopause and sex hormones on the organization of functional brain networks within aging females.

**Key Points:**

- Segregation of functional brain networks declines with increasing age
- Age-segregation relationships are modified by biological sex
- Reproductive stage may impact sex differences in brain network organization

## Introduction

With older age comes normative functional differences in both cognitive and motor domains (Harada et al., 2013; Leal and Yassa, 2014; Stöckel et al., 2017). These age-related behavioral differences are linked to structural differences in brain volume (Raz and Rodrigue, 2006; Bernard and Seidler, 2013), as well as differences in the large-scale organization of brain networks (Chan et al., 2014; Damoiseaux, 2017; King et al., 2018). Thus, understanding the factors that contribute to these brain-behavior relationships is important for advancing care and improving quality of life for the aging population.

Many aging investigations focus on task-based functional activation (Mirelman et al., 2017; Qin and Basak, 2020), though connectivity in the absence of a task is also informative for assessing differences in brain organization over the course of the adult lifespan (Ferreira and Busatto, 2013). The organization of functional brain networks is partially defined by network segregation, which represents greater within-network connectivity strength relative to between-network strength. Network segregation is thought to benefit specialized information processing and efficiency (Bullmore and Sporns, 2012; Wig, 2017) and is often evaluated in comparison to network integration, or dedifferentiation, which corresponds to greater connectivity between networks.

These measures of brain network organization are impacted by age. Young adults demonstrate multiple segregated functional networks with unique behavioral contributions (Power et al., 2011). However, network segregation is typically reduced in advanced age, resulting in increased integration/dedifferentiation of functional brain networks (Goh, 2011; Chan et al., 2014; Geerligs et al., 2015; Setton et al., 2022). Importantly, reduced network segregation, decreased modularity, and dedifferentiation are associated with worsened cognitive and motor performance (King et al., 2018; Kong et al., 2020). In fact, some work shows that network segregation mediates relationships between neurotransmitter systems and behavior in later life (Cassady et al., 2019). As such, the literature suggests that differences in the organization of functional brain networks across adulthood may be a key contributor to age-related behavioral declines.

Notably, aging females are more affected by behavioral and brain differences, compared to males. For example, females demonstrate faster declines in global cognition and greater deficits in balance than males (Wolfson et al., 1994; Levine et al., 2021). Relatedly, females incur higher risk for age-related diseases, such as Alzheimer’s disease (Gao et al., 1998; Alzheimer’s Association, 2021). These sex-specific impacts of age may be related to biological characteristics, such as sex hormone changes with menopause. Menopause is characterized by the cessation of ovarian function, which initiates a decrease in estrogen and progesterone levels. Estrogen and progesterone have been shown to benefit cognition and brain health (Duka et al., 2000; Jacobs and D’Esposito, 2011; Singh and Su, 2013; Hara et al., 2015), and brain circuitry has been associated with hormonal fluctuations across the menstrual cycle (Jacobs et al., 2017; Pritschet et al., 2020). Thus, the loss of neuroprotective hormones with menopause may contribute to disproportionate aging impacts on older females.

However, research on network segregation with respect to sex differences and reproductive aging is lacking. The influence of menopause on the severity of functional declines in aging females is important to factor in when interrogating the origins of sex-specific differences in aging. Such insight would offer important new avenues through which age-related declines may be more effectively addressed. Given the increased incidence and severity of age-related diseases (e.g., Alzheimer’s disease) in females, this knowledge may also promote efforts in the early detection and treatment of disease progression. To address these gaps, we investigated differences in restingstate network segregation between females and males, as well as between reproductive and postmenopausal females, across several functional brain networks. We also looked at associations between network segregation and age across sexes, replicating past research (Chan et al., 2014; Cassady et al., 2019), and within females and males separately.

In the interest of evaluating both cortical and subcortical network segregation, we included ten cortical networks, as defined by Power et al. (2011), and one subcortical network, following Hausman et al. (2020). For the subcortical network, we included striatal seeds originally from Di Martino et al. (2008) and lobular cerebellar seeds that were created using the SUIT atlas (Diedrichsen, 2006; Diedrichsen et al., 2009). Striatal seeds were localized to the left hemisphere while cerebellar seeds were placed in the right hemisphere, given the known lateralization of cerebellar networks with cortical regions. These subcortical regions have reported age differences in connectivity, wherein older adults primarily show reduced resting-state connectivity relative to young adults (Hausman et al., 2020), and are implicated in both motor and cognitive function (Stoodley and Schmahmann, 2009; Stoodley et al., 2012; Helie et al., 2013; Bernard et al., 2017; King et al., 2019). As such, subcortical networks may show differences in brain organization with advanced age, potentially contributing to age-related functional declines. Therefore, to follow-up on prior work and further explore subcortical structures in the context of aging, we included this cerebellar-basal ganglia network as an additional point of comparison in our analyses.

The current investigation was designed to answer several overarching questions. *Question 1:* Can we replicate prior findings showing reduced segregation in cortical networks with increasing age? Given that several studies have shown lower functional network segregation in advanced age (Goh, 2011; Chan et al., 2014; Geerligs et al., 2015; Setton et al., 2022), we predicted that similar age-segregation associations would be present in the current sample. *Question 2:* Do we see the same age-segregation relationships with subcortical structures? Considering the role of the cerebellum and basal ganglia in behaviors associated with age-related declines (Stoodley and Schmahmann, 2009; Stoodley et al., 2012; Helie et al., 2013; Bernard et al., 2017; King et al., 2019), we anticipated that reduced network segregation with increased age would also emerge within this subcortical network. *Question 3:* Do patterns of network segregation declines with age differ between females and males? As females generally experience heavier burdens with older age (Wolfson et al., 1994; Gao et al., 1998; Alzheimer’s Association, 2021; Levine et al., 2021), we predicted that reduced network segregation would be more pronounced in females, relative to males. *Question 4:* Does reproductive stage play a role in potential sex differences with age-segregation relationships? With the benefits of sex hormones in mind (Duka et al., 2000; Jacobs and D’Esposito, 2011; Singh and Su, 2013; Hara et al., 2015), we expected to see greater differences in network segregation between female reproductive stages, as related to hormone loss with menopause, compared to age-matched male controls. These questions are revisited when discussing the present findings.

## Materials and Methods

### Study Sample

Data was accessed through the Cambridge Centre for Ageing and Neuroscience (Cam-CAN) repository (Shafto et al., 2014; Taylor et al., 2017). Data for this repository was gathered from a large sample of healthy adults, ranging from 18 to 88 years of age. We used raw structural and resting-state magnetic resonance imaging (MRI) data, along with demographic variables including sex, age, and menstrual cycle characteristics. We initially acquired data for 652 participants; however, 54 of those participants were excluded for being left-handed or for lacking handedness data. Handedness was assessed using the Edinburgh Handedness Inventory (Oldfield, 1971). This exclusionary criterion was applied to avoid the potential influence of brain organization differences between left-handed and right-handed individuals (Levy and Reid, 1978; Li et al., 2014). Additional individuals were excluded due to MRI data discrepancies, such as missing resting-state scans (n = 4), significant motion artifacts that could not be corrected during image preprocessing (n = 1), and lack of full resting-state volumes (n = 3). As a result, our initial sample consisted of 590 right-handed participants (297 females).

### Reproductive Stage Groupings

Our approach for categorizing females into reproductive, late perimenopausal, early postmenopausal, and late postmenopausal groups was replicated from previous work (Ballard et al., 2021). Here, a brief overview is provided. We used the Stages of Reproductive Aging Workshop Criteria (STRAW+10) to assign females to each reproductive stage group (Harlow et al., 2012). To distinguish between reproductive and late perimenopausal females, we used the reported length of menstrual cycles in days and number of days since last menstrual period. Females with 0-59 days for both variables were classified as reproductive, and females with 60-365 days were put in the late perimenopause group. Further, those within one year of their final menstrual period were also included in the late perimenopause group. To separate postmenopausal females into early and late groups, we used the number of years since final menstrual period. Females with 2-8 years since their final menstrual period were categorized as early postmenopausal, while those with 9+ years since their final menstrual period were assigned to the late postmenopause group.

Females lacking data for menstrual cycle characteristics were categorized by age cut-offs (n = 24): 18-39 for reproductive, 40-49 for late perimenopausal, 55-70 for early postmenopausal, and 71 or older for late postmenopausal. Females ages 50-54, lacking menstrual cycle data, were excluded from final analyses (n = 5) due to variability in reproductive stage for females in this age range (Kato et al., 1998; Morabia and Costanza, 1998; Palmer et al., 2003). To minimize external influences on hormone levels and examine impacts of natural menopause, we excluded females with an intrauterine device (IUD) (n = 12), possible use of continuous birth control (n = 2), and history of hysterectomy (n = 1). Notably, we only excluded females who indicated a hysterectomy that were less than 71 years of age, given that those over the age of 71 with a hysterectomy (n = 7) are likely in a comparable hormonal state to naturally menopausal females of a similar age. The resulting groups from this staging approach were corroborated with subjective responses from females regarding the occurrence of menopause. For further details on our grouping approach, please refer to Ballard et al. (2021).

### Age-Matching

To help account for the intrinsic impact of age on reproductive stage, we formed age-matched male control groups to be used as an indirect reference for female groups. Each male was matched to a female using age, resulting in 1:1 age-matching, along with two variables of quality assurance where necessary: number of outlier scans and maximum motion. Females and males did not significantly differ in either quality assurance variable (*ps* ≥ 0.09). When presented with multiple males of the same age, we chose the male with the number of outlier scans most similar to that of the female in question. If males of the same age also contained identical counts for outlier scans, we chose the male whose maximum motion value was closest to that of the female. In cases where there were more females than males for a particular age, the same approach using number of outlier scans and maximum motion was used to choose female matches. Un-matched males and females were excluded from analyses (n = 156, 70 females); thus, our final sample consisted of 414 participants (207 females, ages 18-87, mean age 56.39 ± 18.80).

Our age-matching method helps account for the natural linkage between age and menopause by facilitating sex comparisons between groups of equal age makeups and sample sizes. In fact, there is notable age overlap between the resulting female groups, even slightly between reproductive (ages 18-55) and late postmenopausal (ages 54-87) females. Characteristics of age-matched groups are reported in **Table 1**. A graphical representation of this data is also available in Ballard et al. (2021), as groups were identical to those used in this prior work.

**Table 1.** Sample characteristics. Sample size and age, in years, for each female reproductive group and relative male control group, after apply age-matching exclusions. The numbers presented here correspond to both the female reproductive group and relative male control group, as 1:1 age-matching resulted in equal age makeups and sample sizes between sexes.

| Stage | Sample Size | Mean Age | Age Range |
| --- | --- | --- | --- |
| Reproductive | 71 | $35.04 \pm 8.80$ | 18-55 |
| Late Perimenopause | 14 | $47.43 \pm 4.70$ | 40-58 |
| Early Postmenopause | 24 | $56.50 \pm 4.91$ | 47-71 |
| Late Postmenopause | 98 | $73.11 \pm 7.74$ | 54-87 |

### Imaging Analyses

A full overview of the study parameters and sample demographics for the Cam-CAN repository can be found in Taylor et al. (2017) and Shafto et al. (2014). For our analyses, we used raw T1 MPRAGE structural scans and raw resting-state EPI scans. The following parameters were used to collect resting-state data: 8 minutes and 30 seconds of acquisition using a 3T Siemens TimTrio, 3 x 3 x 4.4 mm voxel size, and repetition time (TR) of 1.97 seconds.

Image preprocessing and analyses were performed using the CONN toolbox, version 19b (Whitfield-Gabrieli and Nieto-Castanon, 2012). We used the default preprocessing pipeline, which consists of realignment and unwarping with motion correction, centering to (0, 0, 0) coordinates, slice-timing correction, outlier detection using a 95^th^ percentile threshold and the Artifact Rejection Toolbox (ART), segmentation of grey matter, white matter, and cerebrospinal fluid, normalization to MNI space, and spatial smoothing with a 5 mm full width at half-maximum (FWHM) Gaussian kernel. A band-pass filter of 0.008-0.099 Hz was applied to denoise data. The threshold for global-signal z-values was set at 3, while the motion correction threshold was set at 0.5 mm. After being de-spiked during denoising to adhere to the global mean, 6-axis motion data and frame-wise outliers were included as first-level covariates.

MNI coordinates for each cortical node were retrieved from Cassady et al. (2019) (originally derived from Power et al. (2011)). The subcortical network included 20 nodes extracted from Hausman et al. (2020); basal ganglia seeds were originally taken from Di Martino et al. (2008) and cerebellar seeds were determined via the SUIT atlas (Diedrichsen, 2006; Diedrichsen et al., 2009). Combining 214 nodes across 10 cortical networks (Power et al., 2011; Cassady et al., 2019) and 20 subcortical nodes for the cerebellar-basal ganglia network (Diedrichsen, 2006; Di Martino et al., 2008; Diedrichsen et al., 2009; Hausman et al., 2020), our final set of ROIs contained 234 nodes across 11 networks (**Supplementary Table 1**). MNI coordinates for each node were translated to voxel coordinates, which were subsequently used to create spherical seeds with 3.5 mm diameters in FSL (Jenkinson et al., 2012). Seeds were then treated as ROIs, and first-level ROI-to-ROI relationships were evaluated with a bivariate correlation approach.

Next, we looked at group differences between female reproductive stages, as well as age-matched male controls, to investigate relationships with age and effects of sex and reproductive stage on network segregation. Replicating the approach of Chan et al. (2014) and Cassady et al. (2019), network segregation values were determined using **Equation 1** below.

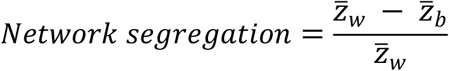

In **Equation 1** (Chan et al., 2014; Cassady et al., 2019), 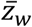 corresponds to the mean correlation between ROIs within an individual network, and 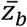 represents the mean correlation between ROIs of an individual network and all remaining ROIs of other networks. Correlation values were transformed into z-values via Fisher’s r-to-z conversion (Zar, 1996). Group-level analyses were performed with a voxel threshold of p < .001 and cluster threshold, FDR-corrected, of p < .05. Imaging analyses for the current work were carried out using the resources provided by the Texas A&M High Performance Research Computing organization.

### Experimental Design and Statistical Analyses

Following Chan et al. (2014) and Cassady et al. (2019), we first investigated relationships between mean network segregation (computed across all networks) and age, as well as age and segregation of each individual network, using Pearson’s correlations. This was carried out with the whole sample. We then completed these analyses within females and males, separately, to elucidate potential sex-specific differences in age-segregation relationships. Finally, to reduce multiple comparisons, we used 2 x 2 between-subjects ANOVAs to evaluate potential effects of sex (females vs. males) and reproductive stage (reproductive vs. late postmenopausal) on mean network segregation and segregation of individual networks.

Consistent with prior work (Chan et al., 2014; Cassady et al., 2019), segregation values above or below three standard deviations from the mean were excluded from final analyses. For analyses considering mean network segregation, exclusions were based on the overall segregation average across all networks, whereas exclusions relative to individual network means were applied to analyses that evaluated each network separately. For the cerebellar-basal ganglia network, one extreme outlier with a segregation value of 697.86, corresponding to 20 standard deviations above the original network mean, was removed. To fairly screen for true outliers in the cerebellar-basal ganglia network, this extreme outlier was removed before performing subsequent exclusions. Values above or below three standard deviations from the adjusted network mean, after removing the extreme outlier, were also excluded as outliers for the cerebellar-basal ganglia network. All statistical analyses were conducted in R programming software.

## Results

### Age Correlations

Detailed results for network segregation and age correlations are reported in **Table 2**. When considering age and mean network segregation (computed across all networks), there is a significant correlation (*p* < 0.001) such that increased age is associated with lower overall network segregation across participants (**Figure 1; Table 2**). Considering each network individually, a significant correlation between age and network segregation emerges for 7 out of the 11 networks investigated: cingulo-opercular task control, default mode, dorsal attention, fronto-parietal task control, salience, sensory somatomotor hand, and visual. Each of these individual network correlations indicate lower network segregation with increased age (**Figure 1; Table 2**).

**Figure 1.**
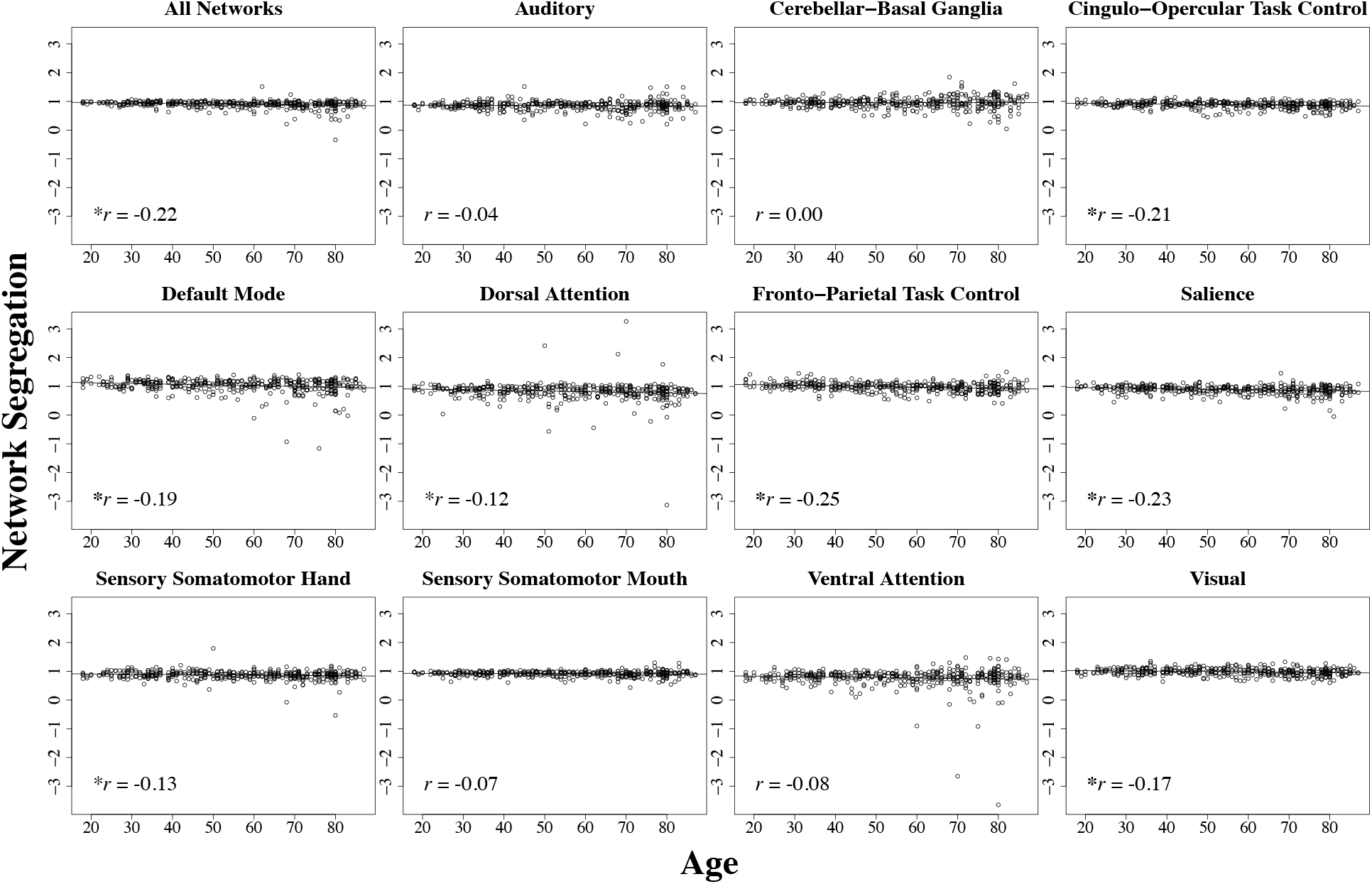
Relationships between age and network segregation across all participants. Distributions for each individual network and the mean across all networks. Each plot includes a linear regression line and the associated correlation value.*: significant p value (at least < .05).

**Table 2.** Pearson’s correlations between age and network segregation across the whole sample. *: p < .05; **:p < .01; ***:p < .001.

| Network | T | Df | P | R |
| --- | --- | --- | --- | --- |
| **Whole Sample** |  |  |  |  |
| All Networks | -4.60 | 408 | ***< 0.001 | -0.22 |
| Auditory | -0.90 | 410 | 0.367 | -0.04 |
| Cerebellar-Basal Ganglia | 0.06 | 408 | 0.950 | 0.00 |
| Cingulo-Opercular Task Control | -4.42 | 405 | ***< 0.001 | -0.21 |
| Default Mode | -3.90 | 410 | ***< 0.001 | -0.19 |
| Dorsal Attention | -2.53 | 411 | *0.012 | -0.12 |
| Fronto-Parietal Task Control | -5.29 | 405 | ***< 0.001 | -0.25 |
| Salience | -4.87 | 408 | ***< 0.001 | -0.23 |
| Sensory Somatomotor Hand | -2.58 | 411 | *0.010 | -0.13 |
| Sensory Somatomotor Mouth | -1.36 | 406 | 0.175 | -0.07 |
| Ventral Attention | -1.71 | 410 | 0.089 | -0.08 |
| Visual | -3.55 | 407 | ***< 0.001 | -0.17 |

Correlations in females only reveal that older females exhibit lower segregation in almost all of the same networks as the whole sample: cingulo-opercular task control, default mode, dorsal attention, fronto-parietal task control, salience, and visual (**Figure 2; Table 3**). Further, females show a significant negative relationship between age and mean network segregation (*p* < 0.001). In contrast to the whole sample results, females do not demonstrate a significant correlation between age and segregation of the sensory somatomotor hand network (*p* = 0.124).

**Figure 2.**
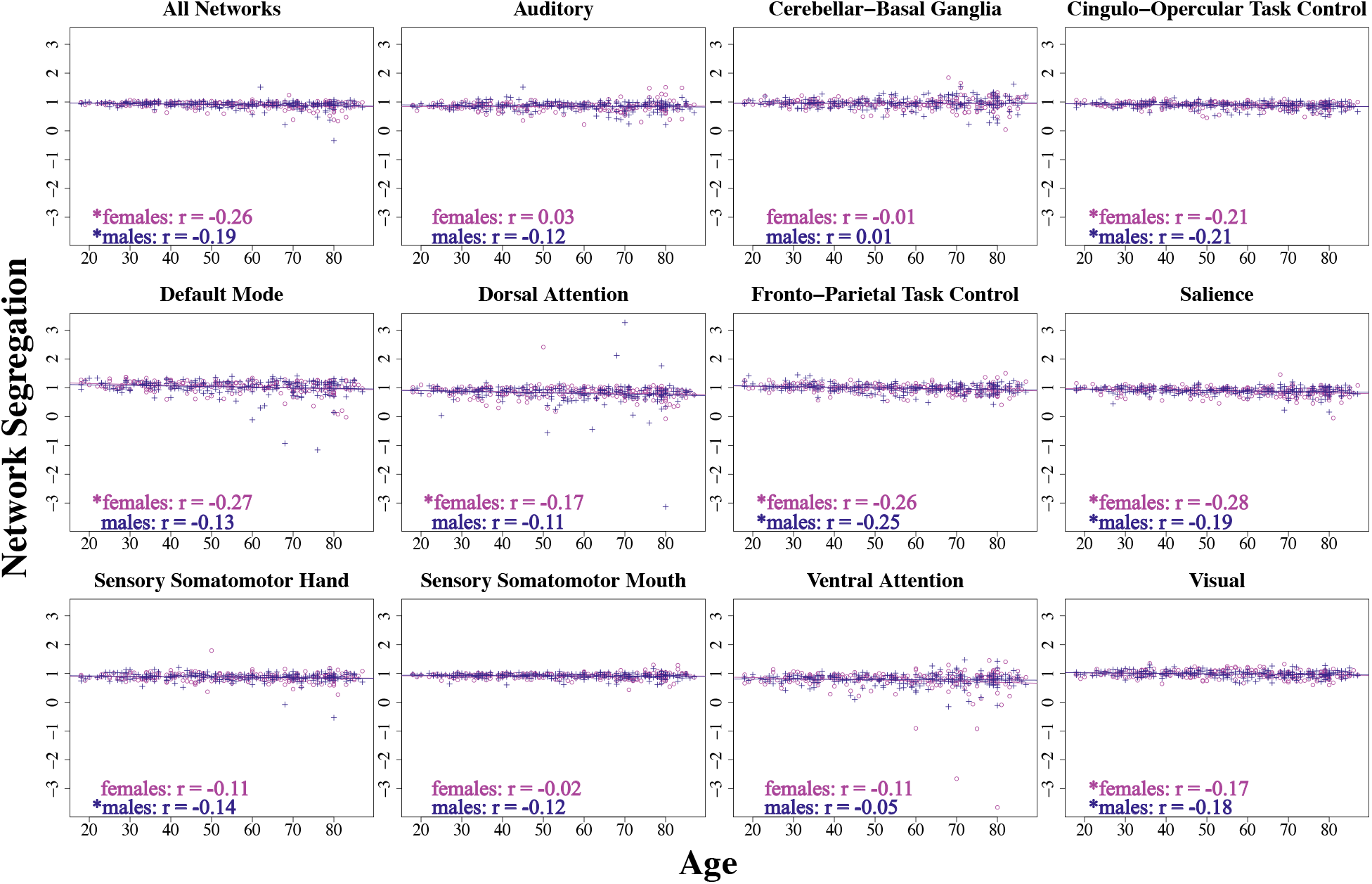
Relationships between age and network segregation, separated by sex. Distributions for each individual network and the mean across all networks. Females are plotted in purple and males in blue. Each plot includes linear regression lines and associated correlation values. *: significant p value (at least < .05).

**Table 3.** Pearson’s correlations between age and network segregation within each sex separately. *:p < .05; **: p < .01; ***: p < .001.

| Network | T | Df | P | R |
| --- | --- | --- | --- | --- |
| **Females Only** |  |  |  |  |
| All Networks | -3.84 | 202 | ***$< 0.001$ | -0.26 |
| Auditory | 0.39 | 203 | 0.694 | 0.03 |
| Cerebellar-Basal Ganglia | -0.09 | 204 | 0.925 | -0.01 |
| Cingulo-Opercular Task Control | -3.13 | 202 | **0.002 | -0.21 |
| Default Mode | -4.03 | 203 | ***$< 0.001$ | -0.27 |
| Dorsal Attention | -2.44 | 204 | *0.016 | -0.17 |
| Fronto-Parietal Task Control | -3.88 | 202 | ***< 0.001 | -0.26 |
| Salience | -4.12 | 203 | ***< 0.001 | -0.28 |
| Sensory Somatomotor Hand | -1.54 | 205 | 0.124 | -0.11 |
| Sensory Somatomotor Mouth | -0.28 | 202 | 0.779 | -0.02 |
| Ventral Attention | -1.58 | 204 | 0.116 | -0.11 |
| Visual | -2.50 | 202 | *0.013 | -0.17 |
| **Males Only** |  |  |  |  |
| All Networks | -2.78 | 204 | **0.006 | -0.19 |
| Auditory | -1.78 | 205 | 0.077 | -0.12 |
| Cerebellar-Basal Ganglia | 0.19 | 202 | 0.853 | 0.01 |
| Cingulo-Opercular Task Control | -3.11 | 201 | **0.002 | -0.21 |
| Default Mode | -1.91 | 205 | 0.058 | -0.13 |
| Dorsal Attention | -1.58 | 205 | 0.116 | -0.11 |
| Fronto-Parietal Task Control | -3.59 | 201 | ***< 0.001 | -0.25 |
| Salience | -2.70 | 203 | **0.007 | -0.19 |
| Sensory Somatomotor Hand | -2.08 | 204 | *0.038 | -0.14 |
| Sensory Somatomotor Mouth | -1.76 | 202 | 0.079 | -0.12 |
| Ventral Attention | -0.66 | 204 | 0.510 | -0.05 |
| Visual | -2.54 | 203 | *0.012 | -0.18 |

In age-matched male controls, we see broadly similar associations between age and network segregation, though some differences emerge as well (**Figure 2; Table 3**). Mirroring females, male controls demonstrate lower segregation with increased age across all networks as well as within the cingulo-opercular task control, fronto-parietal task control, salience, and visual networks. However, unique to males, negative associations between age and network segregation are also present for the sensory somatomotor hand network (*p* = 0.038). Interestingly, males do not demonstrate significant correlations between age and network segregation for the default mode and dorsal attention networks; thus, those relationships are unique to females. Overall, it seems that though some similarities exist between females and males, a few sex-specific differences in functional network segregation declines with age are present in the current sample.

### Reproductive Stage Comparisons

An overview of reproductive stage and sex effects on network segregation is provided in **Table 4**. An effect of stage (reproductive vs. late postmenopausal/relative controls) was significant across networks when considered together (*p* < 0.001) as well as within 6 individual networks: cingulo-opercular task control, default mode, fronto-parietal task control, salience, sensory somatomotor hand, and visual. Notably, however, the effect of sex (females vs. males) was not statistically significant across networks, both on average and for the individual networks considered here. In addition, interactions between reproductive stage and sex were not significant for any networks. As such, given the lack of sex effects or interactions between stage and sex, this series of analyses suggests that reproductive stage within females does not differentially impact functional network segregation beyond the impacts of age more generally. The consistent significance of main effects for reproductive stage across both sexes may be more directly attributed to age alone (**Figure 3**). However, exploratory analyses with the reproductive and early postmenopause groups indicate that sex-specific differences in network segregation, with respect to reproductive stage, are present during the transition to menopause (see **Supplementary Table 2**). This may suggest that the menopausal transition is particularly important for network dynamics, but in later life once hormones reach a more stable low state, sex differences are no longer present. However, we would note that this is exploratory and speculative.

**Figure 3.**
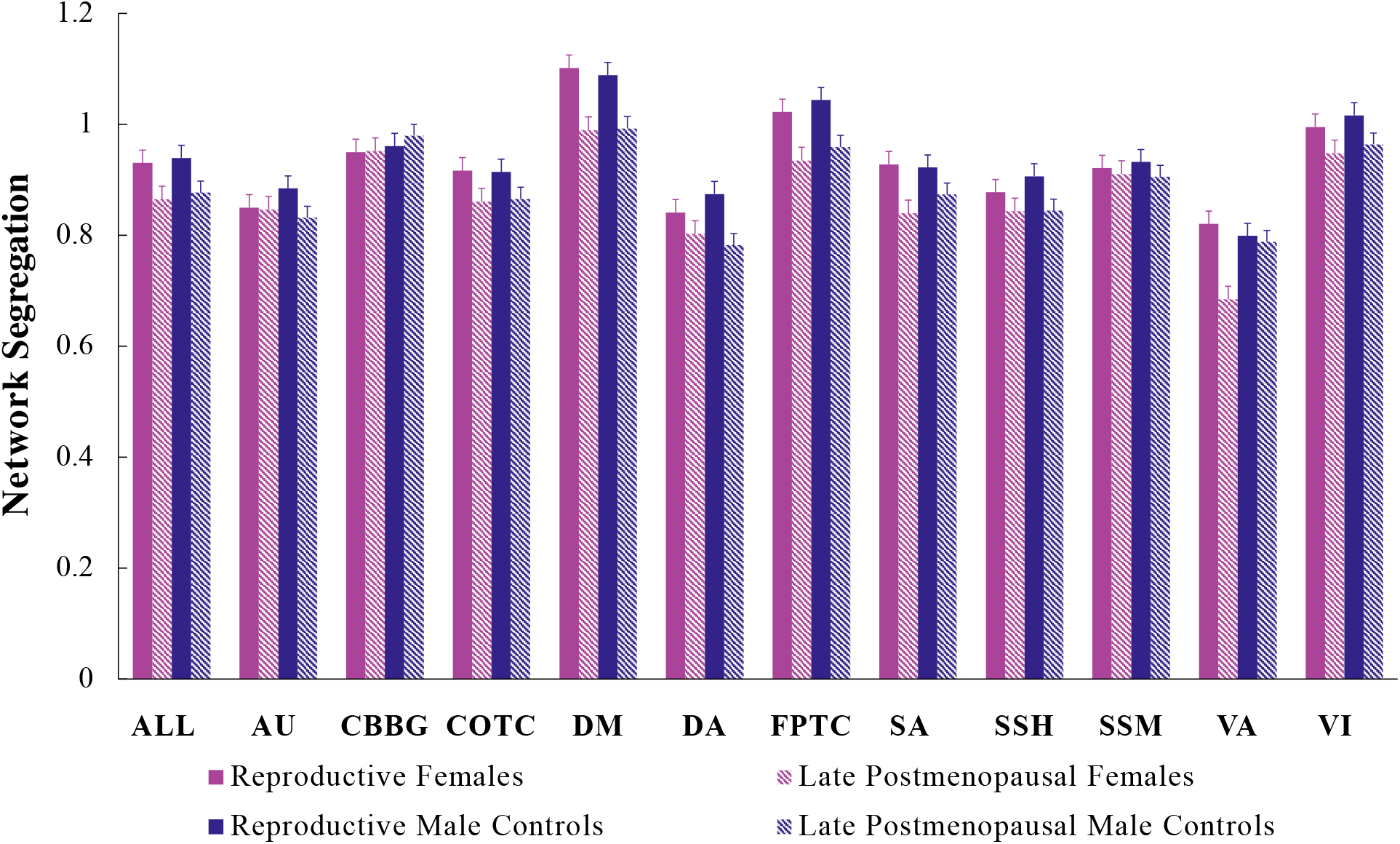
Network segregation by stage (reproductive vs. late postmenopausal/relative controls) and sex (female vs. male). Average segregation value per group for individual networks and the mean across all networks. Error bars depict standard error. ALL: Mean across all networks; AU: Auditory; CBBG: Cerebellar-basal ganglia; COTC: Cingulo-opercular task control; DM: Default mode; DA: Dorsal attention; FPTC: Fronto-parietal task control; SA: Salience; SSH: Sensory somatomotor hand; SSM: Sensory somatomotor mouth; VA: Ventral attention; VI: Visual.

**Table 4.**
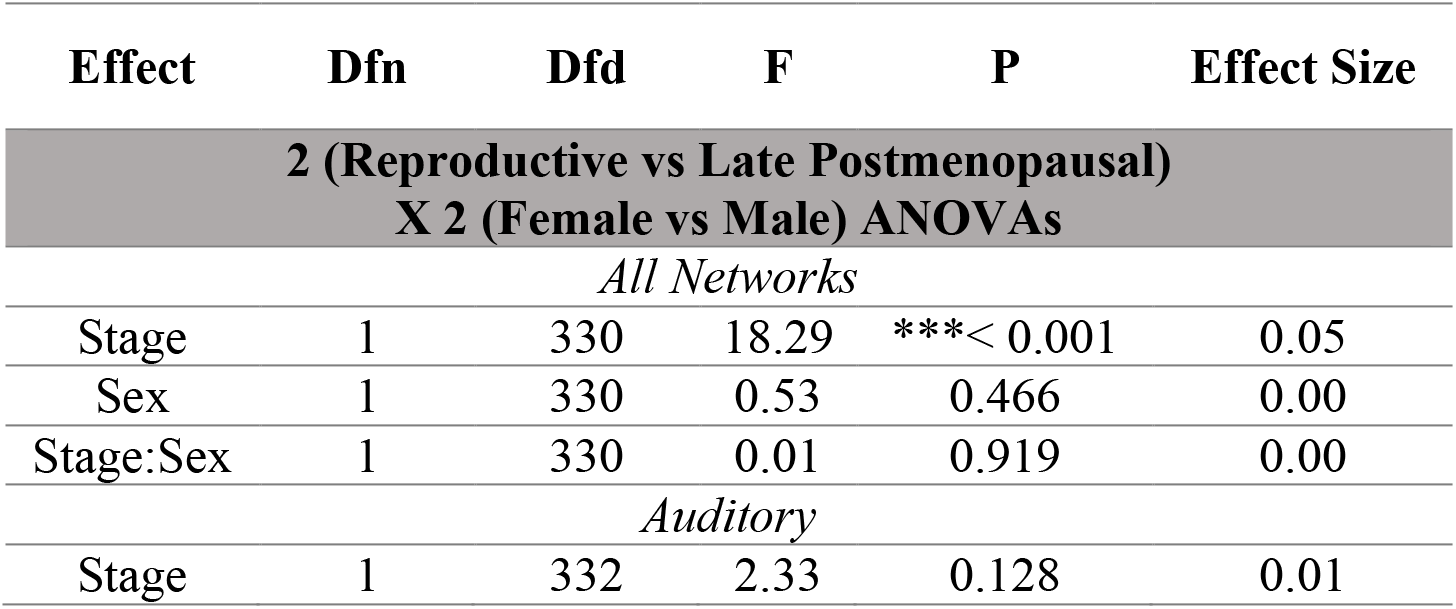
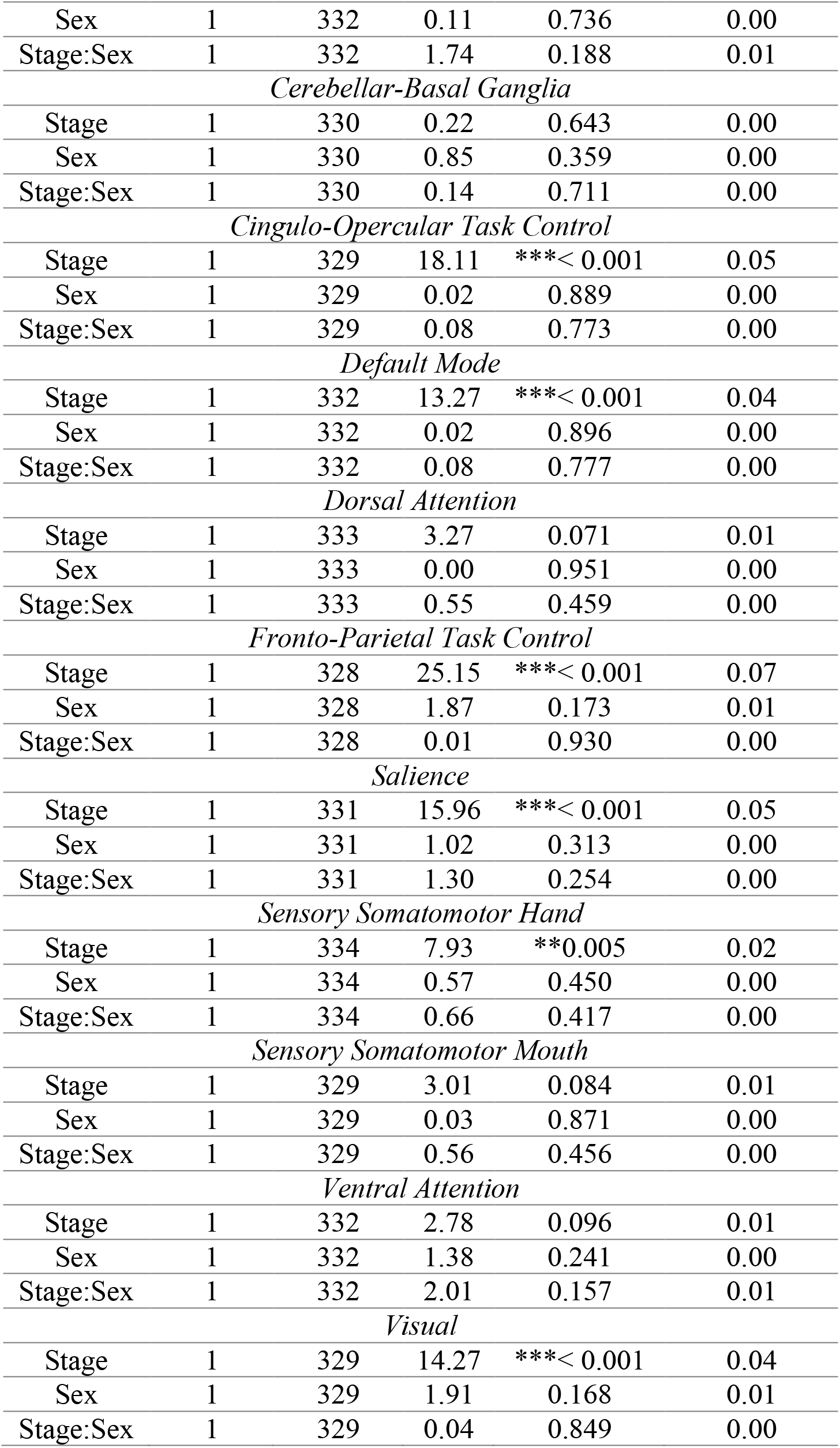
Between-subjects ANOVA results. *:p < .05; **: p < .01; ***:p < .001.

## Discussion

In the current study, we investigated age-related differences in functional network segregation in the context of sex and female reproductive stage. Following previous work (Chan et al., 2014; Cassady et al., 2019), we examined ten cortical networks (Power et al., 2011) and also included an additional subcortical network (Diedrichsen, 2006; Di Martino et al., 2008; Diedrichsen et al., 2009; Hausman et al., 2020). We first replicated past work showing lower network segregation with higher age across participants, as well as within females and males separately, answering *Question 1* and consistent with our predictions. Contrary to what we predicted for *Question 2* however, we did not find significant age-segregation relationships with the subcortical network. Next, when evaluating relationships between age and network segregation within each sex separately, we found distinct patterns between females and males, along with some similarities, which supports our predictions related to *Question 3*. Finally, we explored effects of sex and reproductive stage on network segregation; these tests revealed significant effects of reproductive stage on functional network segregation, while effects of sex and stage by sex interactions were null. These particular results leave *Question 4* unanswered, as more work is needed to define the role of menopause in sex differences with age-related network segregation declines. Results are further discussed in the context of the relevant literature below.

### Resting-state network segregation and aging

When looking at associations between mean network segregation and age across all participants, we find a significant correlation in the negative direction, indicating lower overall network segregation with increased age. This parallels the overarching theme in the literature, wherein older age is associated with lower network segregation and, in turn, greater integration between networks (Chan et al., 2014; Geerligs et al., 2015; Grady et al., 2016; Spreng et al., 2016; King et al., 2018; Cassady et al., 2019). As functional network segregation is associated with specialized information processing and efficiency (Bullmore and Sporns, 2012), as well as successful cognition and motor function (King et al., 2018; Kong et al., 2020), age-related differences in network segregation, as observed here and in prior work, may contribute to functional declines in the aging population. This shift from network segregation to integration, or dedifferentiation, may represent a compensatory mechanism for the natural depreciation of brain function with age, in turn, impacting functional performance.

Relatedly, when investigating age-segregation relationships within individual networks (using the whole sample), we found that segregation was lower with higher age in 7 out of 11 functional networks: cingulo-opercular task control, default mode, dorsal attention, fronto-parietal task control, salience, sensory somatomotor hand, and visual. This is broadly consistent with findings from Chan et al. (2014) and Cassady et al. (2019), who reported similar relationships in 8/10 and 5/10 networks, respectively. We observed age-related relationships with network segregation in regions responsible for several behavioral domains, such as adaptive task control, spontaneous cognition, top-down attentional control, and working memory (Andrews-Hanna et al., 2010; Vossel et al., 2014; Wallis et al., 2015; Marek and Dosenbach, 2018).

We did not observe significant correlations with age in 4 of the 11 networks: auditory, cerebellar-basal ganglia, sensory somatomotor mouth, and ventral attention. This indicates that, contrary to our expectations, subcortical network segregation, at least in the cerebellar-basal ganglia network, is not correlated with age. Notably, we investigated this network as a whole for the current investigation, though in our prior work, we did show some degree of functional dedifferentiation *within* the cerebellar-basal ganglia network in older adults. That is, motor nodes became less strongly associated with one another as did nodes associated with cognitive networks and structures (Hausman et al., 2020). Here, in relation to other cortical networks, we did not see any age-segregation associations for this specific subcortical network. As such, we suggest that within network dynamics change and show evidence for dedifferentiation subcortically, but this is distinct from broader global dynamics with cortical networks.

In addition, this demonstrates that networks responsible for attentional filtering (ventral attention) (Vossel et al., 2014) and the interpretation of sensory information (auditory and sensory somatomotor hand) (Kayser et al., 2005; Small and Green, 2012) do not show the same lessened segregation in older age as the other networks investigated. However, this is unlike Cassady et al. (2019) who found significant correlations between functional segregation and age in the auditory and sensory somatomotor mouth networks, whereas we did not. Moreover, Chan et al. (2014) did not find associations with age in the sensory somatomotor hand and salience networks, while such relationships were in fact observed here. Importantly, our study included 414 participants, whereas previous works had substantially smaller samples, which may partially explain some of the observed differences between our results and findings from the larger body of work on network segregation in older adulthood. However, the mixed findings across these studies collectively suggest that the relationships between segregation of these networks and age are certainly less reliable and robust than other cortical networks.

### Sex-specific differences in network segregation

When breaking down age and network segregation associations by sex, we find both similarities and differences in females and males. Both sexes exhibit lower segregation with older age in cingulo-opercular task control, fronto-parietal task control, salience, and visual networks, though females also demonstrate negative correlations with age in the default mode and dorsal attention networks while these relationships were not seen in males. Thus, females may endure greater consequences with age in respect to the organization and efficiency of these particular functional networks, which may contribute to the disproportionate impact of normative behavioral declines and age-related disease on aging females, compared to males. Interestingly, the default mode network is strongly implicated in Alzheimer’s disease pathology, which is more prevalent and severe in females (Greicius et al., 2004; Jones et al., 2011; Alzheimer’s Association, 2021). Suppression of default mode network activity is associated with better performance on cognitive tasks (Anticevic et al., 2012). Consequently, the lack of default mode network segregation with age, specifically in females, may also reflect an inability to successfully inhibit default mode function during task-positive processing, in turn, contributing to age-related behavioral deficits.

On the other hand, males demonstrate unique age-related declines in segregation of the sensory somatomotor hand network, whereas females do not. Interestingly, in our prior work examining resting-state connectivity differences in cerebellar-whole brain networks between female reproductive stages and age-matched males, we found that male control groups exhibited greater differences in cerebellar-somatosensory connectivity compared to female reproductive groups (Ballard et al., 2021). Specifically, late postmenopausal male controls showed lower connectivity between cerebellar regions associated with cognition (Crus I/II) and regions of the somatosensory cortex, relative to both reproductive male controls and female counterparts. Though this prior work used the same sample from the current investigation, our results here align with these previous findings and illustrate that connectivity differences in somatosensory regions, specific to males, stand when using an alternative analysis approach. In sum, our findings highlight that sex-specific differences are important to consider when exploring relationships between age and the organization of functional brain networks.

Moreover, comparisons between female reproductive stages and age-matched male controls offer additional insight on a possible link between sex and network segregation differences. Notably, in our ANOVAs, reproductive stage influenced segregation for several networks while no effects of sex were observed. These findings may point to age effects, though impacts of sex hormone changes with menopause may also be at play and would be useful for better categorizing reproductive stage. Notably, in an exploratory analysis evaluating network segregation differences between reproductive and early postmenopausal groups, effects of sex and interactions between reproductive stage and sex begin to emerge in a few networks (**Supplementary Table 2**). Though these results are not conclusive, this indicates that the transition to menopause and initial declines in sex hormones may be important to evaluate in the context of sex differences in aging outcomes, however after the menopausal transition there are fewer sex differences. As such, it may be the case that the dynamics of hormonal change are important in midlife, though we would caution that this is highly speculative at this point. We would also note that, without access to direct hormone data we cannot accurately tease out these potential influences on the present results. Further work assessing direct effects of sex hormones is needed to fully understand the impact of menopause on the brain in aging females.

To this point, the current body of work on network segregation in older age does not include sex-specific analyses or comparisons between female reproductive stages (Chan et al., 2014; Geerligs et al., 2015; Grady et al., 2016; Spreng et al., 2016; King et al., 2018; Cassady et al., 2019). In fact, none of this work reports any analyses on network segregation differences between females and males. However, females endure more severe functional declines with age (Wolfson et al., 1994; Levine et al., 2021); thus, functional network segregation differences may contribute, at least in part, to the imbalance in aging trajectories between females and males. Results from the present work suggest that sex is a crucial consideration when examining the organization of resting-state networks with respect to aging, and more work including the potential influence of sex steroid hormones is needed in the context of female reproductive aging. Future investigations should include sex-specific analyses and evaluate the effects of hormone changes with menopause when investigating brain differences in older adulthood.

### Limitations

Though this investigation contributes to current advances in aging research, there are limitations worth noting. First, we lacked access to direct hormone data or data regarding consecutive cycle lengths for our reproductive stage categorizations. As a result, reproductive stage was characterized using self-report menstrual information, and females undergoing hormone therapy or taking hormonal contraceptives may have been included in our sample. As the effects of menopause on functional network segregation may be more explicitly linked to hormone fluctuations, as opposed to broad reproductive stage differences, the lack of hormone data has limited our investigation. Moreover, given the lack of consecutive cycle data, our reproductive group may inherently include females in early perimenopause. Second, we did not evaluate behavioral performance. Therefore, interpretation of functional relevance is purely theoretical. Notably, we did not correct for multiple comparisons in our analyses and results should be interpreted with caution, though this approach follows the previous work from Chan et al. (2014) and Cassady et al. (2019), and we sought to replicate their approaches as closely as possible. This includes our approach relative to multiple comparisons correction. Finally, given that menopause is a product of aging, we cannot discount impacts of age on the current findings. However, female reproductive groups overlap in age and age-matched male controls help to limit age impacts.

## Conclusion

The current study, using data from the CamCAN repository, offers new insight into sexspecific differences in the aging brain. Here, we evaluated the influence of sex and female reproductive stage on age-related associations with functional network segregation. We provide evidence for distinct patterns of functional network segregation between females and males, along with potential effects of reproductive stage, indicating that these biological factors may contribute to some degree to the differing aging trajectories between sexes. However, subsequent work is needed to determine the particular role of sex hormone fluctuations with menopause on brain differences within aging females. Such work is necessary to support findings from the present investigation and provide potential avenues through which age-related declines may be alleviated. Further, given sex differences in non-normative aging, elucidating relationships between menopause and the aging brain may also offer treatment alternatives for age-related diseases, such as Alzheimer’s disease and other dementias.

## Conflict of Interest

The authors declare no competing financial or alternative interests.

## Acknowledgements

Data for the current investigation was accessed through the Cambridge Centre for Ageing and Neuroscience (Cam-CAM) repository. Funding for the Cam-CAN repository was provided by the UK Biotechnology and Biological Sciences Research Council (grant number BB/H008217/1), along with the University of Cambridge, UK and the UK Medical Research Council. The current study was also supported by funding from the National Institute on Aging to JAB (grant number R01AG065010).

## Data Availability

All data incorporated in the present study was accessed through the Cam-CAN online repository and is available for shared use at https://camcan-archive.mrc-cbu.cam.ac.uk/dataaccess/.

## Notes

https://camcan-archive.mrc-cbu.cam.ac.uk/dataaccess/

### Competing Interest Statement

The authors have declared no competing interest.

