## Supplemental Tables for "Age-related differences in functional network segregation in the context of sex and reproductive stage"

**Supplementary Tables**

| Network | ROI # | MNI Coordinates | | |
| --- | --- | --- | --- | --- |
|  |  | **X** | **Y** | **Z** |
| Sensory Somatomotor Hand | 1 | -7 | -52 | 61 |
| Sensory Somatomotor Hand | 2 | -14 | -18 | 40 |
| Sensory Somatomotor Hand | 3 | 0 | -15 | 47 |
| Sensory Somatomotor Hand | 4 | 10 | -2 | 45 |
| Sensory Somatomotor Hand | 5 | -7 | -21 | 65 |
| Sensory Somatomotor Hand | 6 | -7 | -33 | 72 |
| Sensory Somatomotor Hand | 7 | 13 | -33 | 75 |
| Sensory Somatomotor Hand | 8 | -54 | -23 | 43 |
| Sensory Somatomotor Hand | 9 | 29 | -17 | 71 |
| Sensory Somatomotor Hand | 10 | 10 | -46 | 73 |
| Sensory Somatomotor Hand | 11 | -23 | -30 | 72 |
| Sensory Somatomotor Hand | 12 | -40 | -19 | 54 |
| Sensory Somatomotor Hand | 13 | 29 | -39 | 59 |
| Sensory Somatomotor Hand | 14 | 50 | -20 | 42 |
| Sensory Somatomotor Hand | 15 | -38 | -27 | 69 |
| Sensory Somatomotor Hand | 16 | 20 | -29 | 60 |
| Sensory Somatomotor Hand | 17 | 44 | -8 | 57 |
| Sensory Somatomotor Hand | 18 | -29 | -43 | 61 |
| Sensory Somatomotor Hand | 19 | 10 | -17 | 74 |
| Sensory Somatomotor Hand | 20 | 22 | -42 | 69 |
| Sensory Somatomotor Hand | 21 | -45 | -32 | 47 |
| Sensory Somatomotor Hand | 22 | -21 | -31 | 61 |
| Sensory Somatomotor Hand | 23 | -13 | -17 | 75 |
| Sensory Somatomotor Hand | 24 | 42 | -20 | 55 |
| Sensory Somatomotor Hand | 25 | -38 | -15 | 69 |
| Sensory Somatomotor Hand | 26 | -16 | -46 | 73 |
| Sensory Somatomotor Hand | 27 | 2 | -28 | 60 |
| Sensory Somatomotor Hand | 28 | 3 | -17 | 58 |
| Sensory Somatomotor Hand | 29 | 38 | -17 | 45 |
| Sensory Somatomotor Hand | 30 | 47 | -30 | 49 |
| Visual | 31 | 18 | -47 | -10 |
| Visual | 32 | 40 | -72 | 14 |
| Visual | 33 | 8 | -72 | 11 |
| Visual | 34 | -8 | -81 | 7 |
| Visual | 35 | -28 | -79 | 19 |
| Visual | 36 | 20 | -66 | 2 |
| Visual | 37 | -24 | -91 | 19 |
| Visual | 38 | 27 | -59 | -9 |
| Visual | 39 | -15 | -72 | -8 |
| Visual | 40 | -18 | -68 | 5 |
| Visual | 41 | 43 | -78 | -12 |
| Visual | 42 | -47 | -76 | -10 |
| Visual | 43 | -14 | -91 | 31 |
| Visual | 44 | 15 | -87 | 37 |
| Visual | 45 | 29 | -77 | 25 |
| Visual | 46 | 20 | -86 | -2 |
| Visual | 47 | 15 | -77 | 31 |
| Visual | 48 | -16 | -52 | -1 |
| Visual | 49 | 42 | -66 | -8 |
| Visual | 50 | 24 | -87 | 24 |
| Visual | 51 | 6 | -72 | 24 |
| Visual | 52 | -42 | -74 | 0 |
| Visual | 53 | 26 | -79 | -16 |
| Visual | 54 | -16 | -77 | 34 |
| Visual | 55 | -3 | -81 | 21 |
| Visual | 56 | -40 | -88 | -6 |
| Visual | 57 | 37 | -84 | 13 |
| Visual | 58 | 6 | -81 | 6 |
| Visual | 59 | -26 | -90 | 3 |
| Visual | 60 | -33 | -79 | -13 |
| Visual | 61 | 37 | -81 | 1 |
| Sensory Somatomotor Mouth | 62 | -49 | -11 | 35 |
| Sensory Somatomotor Mouth | 63 | 36 | -9 | 14 |
| Sensory Somatomotor Mouth | 64 | 51 | -6 | 32 |
| Sensory Somatomotor Mouth | 65 | -53 | -10 | 24 |
| Sensory Somatomotor Mouth | 66 | 66 | -8 | 25 |
| Auditory | 67 | 32 | -26 | 13 |
| Auditory | 68 | 65 | -33 | 20 |
| Auditory | 69 | 58 | -16 | 7 |
| Auditory | 70 | -38 | -33 | 17 |
| Auditory | 71 | -60 | -25 | 14 |
| Auditory | 72 | -49 | -26 | 5 |
| Auditory | 73 | 43 | -23 | 20 |
| Auditory | 74 | -50 | -34 | 26 |
| Auditory | 75 | -53 | -22 | 23 |
| Auditory | 76 | -55 | -9 | 12 |
| Auditory | 77 | 56 | -5 | 13 |
| Auditory | 78 | 59 | -17 | 29 |
| Auditory | 79 | -30 | -27 | 12 |
| Default Mode | 80 | -41 | -75 | 26 |
| Default Mode | 81 | 6 | 67 | -4 |
| Default Mode | 82 | 8 | 48 | -15 |
| Default Mode | 83 | -13 | -40 | 1 |
| Default Mode | 84 | -18 | 63 | -9 |
| Default Mode | 85 | -46 | -61 | 21 |
| Default Mode | 86 | 43 | -72 | 28 |
| Default Mode | 87 | -44 | 12 | -34 |
| Default Mode | 88 | 46 | 16 | -30 |
| Default Mode | 89 | -68 | -23 | -16 |
| Default Mode | 90 | -44 | -65 | 35 |
| Default Mode | 91 | -39 | -75 | 44 |
| Default Mode | 92 | -7 | -55 | 27 |
| Default Mode | 93 | 6 | -59 | 35 |
| Default Mode | 94 | -11 | -56 | 16 |
| Default Mode | 95 | -3 | -49 | 13 |
| Default Mode | 96 | 8 | -48 | 31 |
| Default Mode | 97 | 15 | -63 | 26 |
| Default Mode | 98 | -2 | -37 | 44 |
| Default Mode | 99 | 11 | -54 | 17 |
| Default Mode | 100 | 52 | -59 | 36 |
| Default Mode | 101 | 23 | 33 | 48 |
| Default Mode | 102 | -10 | 39 | 52 |
| Default Mode | 103 | -16 | 29 | 53 |
| Default Mode | 104 | -35 | 20 | 51 |
| Default Mode | 105 | 22 | 39 | 39 |
| Default Mode | 106 | 13 | 55 | 38 |
| Default Mode | 107 | -10 | 55 | 39 |
| Default Mode | 108 | -20 | 45 | 39 |
| Default Mode | 109 | 6 | 54 | 16 |
| Default Mode | 110 | 6 | 64 | 22 |
| Default Mode | 111 | -7 | 51 | -1 |
| Default Mode | 112 | 9 | 54 | 3 |
| Default Mode | 113 | -3 | 44 | -9 |
| Default Mode | 114 | 8 | 42 | -5 |
| Default Mode | 115 | -11 | 45 | 8 |
| Default Mode | 116 | -2 | 38 | 36 |
| Default Mode | 117 | -3 | 42 | 16 |
| Default Mode | 118 | -20 | 64 | 19 |
| Default Mode | 119 | -8 | 48 | 23 |
| Default Mode | 120 | 65 | -12 | -19 |
| Default Mode | 121 | -56 | -13 | -10 |
| Default Mode | 122 | -58 | -30 | -4 |
| Default Mode | 123 | 65 | -31 | -9 |
| Default Mode | 124 | -68 | -41 | -5 |
| Default Mode | 125 | 13 | 30 | 59 |
| Default Mode | 126 | 12 | 36 | 20 |
| Default Mode | 127 | 52 | -2 | -16 |
| Default Mode | 128 | -26 | -40 | -8 |
| Default Mode | 129 | 27 | -37 | -13 |
| Default Mode | 130 | -34 | -38 | -16 |
| Default Mode | 131 | 28 | -77 | -32 |
| Default Mode | 132 | 52 | 7 | -30 |
| Default Mode | 133 | -53 | 3 | -27 |
| Default Mode | 134 | 47 | -50 | 29 |
| Default Mode | 135 | -49 | -42 | 1 |
| Default Mode | 136 | -46 | 31 | -13 |
| Default Mode | 137 | 49 | 35 | -12 |
| Fronto-Parietal Task Control | 138 | -44 | 2 | 46 |
| Fronto-Parietal Task Control | 139 | 48 | 25 | 27 |
| Fronto-Parietal Task Control | 140 | -47 | 11 | 23 |
| Fronto-Parietal Task Control | 141 | -53 | -49 | 43 |
| Fronto-Parietal Task Control | 142 | -23 | 11 | 64 |
| Fronto-Parietal Task Control | 143 | 58 | -53 | -14 |
| Fronto-Parietal Task Control | 144 | 24 | 45 | -15 |
| Fronto-Parietal Task Control | 145 | 34 | 54 | -13 |
| Fronto-Parietal Task Control | 146 | 47 | 10 | 33 |
| Fronto-Parietal Task Control | 147 | -41 | 6 | 33 |
| Fronto-Parietal Task Control | 148 | -42 | 38 | 21 |
| Fronto-Parietal Task Control | 149 | 38 | 43 | 15 |
| Fronto-Parietal Task Control | 150 | 49 | -42 | 45 |
| Fronto-Parietal Task Control | 151 | -28 | -58 | 48 |
| Fronto-Parietal Task Control | 152 | 44 | -53 | 47 |
| Fronto-Parietal Task Control | 153 | 32 | 14 | 56 |
| Fronto-Parietal Task Control | 154 | 37 | -65 | 40 |
| Fronto-Parietal Task Control | 155 | -42 | -55 | 45 |
| Fronto-Parietal Task Control | 156 | 40 | 18 | 40 |
| Fronto-Parietal Task Control | 157 | -34 | 55 | 4 |
| Fronto-Parietal Task Control | 158 | -42 | 45 | -2 |
| Fronto-Parietal Task Control | 159 | 33 | -53 | 44 |
| Fronto-Parietal Task Control | 160 | 43 | 49 | -2 |
| Fronto-Parietal Task Control | 161 | -42 | 25 | 30 |
| Fronto-Parietal Task Control | 162 | -3 | 26 | 44 |
| Ventral Attention | 163 | -10 | 11 | 67 |
| Ventral Attention | 164 | 54 | -43 | 22 |
| Ventral Attention | 165 | -56 | -50 | 10 |
| Ventral Attention | 166 | -55 | -40 | 14 |
| Ventral Attention | 167 | 52 | -33 | 8 |
| Ventral Attention | 168 | 51 | -29 | -4 |
| Ventral Attention | 169 | 56 | -46 | 11 |
| Ventral Attention | 170 | 53 | 33 | 1 |
| Ventral Attention | 171 | -49 | 25 | -1 |
| Cingulo-Opercular Task Control | 172 | -3 | 2 | 53 |
| Cingulo-Opercular Task Control | 173 | 54 | -28 | 34 |
| Cingulo-Opercular Task Control | 174 | 19 | -8 | 64 |
| Cingulo-Opercular Task Control | 175 | -16 | -5 | 71 |
| Cingulo-Opercular Task Control | 176 | -10 | -2 | 42 |
| Cingulo-Opercular Task Control | 177 | 37 | 1 | -4 |
| Cingulo-Opercular Task Control | 178 | 13 | -1 | 70 |
| Cingulo-Opercular Task Control | 179 | 7 | 8 | 51 |
| Cingulo-Opercular Task Control | 180 | -45 | 0 | 9 |
| Cingulo-Opercular Task Control | 181 | 49 | 8 | -1 |
| Cingulo-Opercular Task Control | 182 | -34 | 3 | 4 |
| Cingulo-Opercular Task Control | 183 | -51 | 8 | -2 |
| Cingulo-Opercular Task Control | 184 | -5 | 18 | 34 |
| Cingulo-Opercular Task Control | 185 | 36 | 10 | 1 |
| Dorsal Attention | 186 | 10 | -62 | 61 |
| Dorsal Attention | 187 | -52 | -63 | 5 |
| Dorsal Attention | 188 | 22 | -65 | 48 |
| Dorsal Attention | 189 | 46 | -59 | 4 |
| Dorsal Attention | 190 | 25 | -58 | 60 |
| Dorsal Attention | 191 | -33 | -46 | 47 |
| Dorsal Attention | 192 | -27 | -71 | 37 |
| Dorsal Attention | 193 | -32 | -1 | 54 |
| Dorsal Attention | 194 | -42 | -60 | -9 |
| Dorsal Attention | 195 | -17 | -59 | 64 |
| Dorsal Attention | 196 | 29 | -5 | 54 |
| Salience | 197 | 11 | -39 | 50 |
| Salience | 198 | 55 | -45 | 37 |
| Salience | 199 | 42 | 0 | 47 |
| Salience | 200 | 31 | 33 | 26 |
| Salience | 201 | 48 | 22 | 10 |
| Salience | 202 | -35 | 20 | 0 |
| Salience | 203 | 36 | 22 | 3 |
| Salience | 204 | 37 | 32 | -2 |
| Salience | 205 | 34 | 16 | -8 |
| Salience | 206 | -11 | 26 | 25 |
| Salience | 207 | -1 | 15 | 44 |
| Salience | 208 | -28 | 52 | 21 |
| Salience | 209 | 0 | 30 | 27 |
| Salience | 210 | 5 | 23 | 37 |
| Salience | 211 | 10 | 22 | 27 |
| Salience | 212 | 31 | 56 | 14 |
| Salience | 213 | 26 | 50 | 27 |
| Salience | 214 | -39 | 51 | 17 |
| Cerebellar-Basal Ganglia | 215 | -47 | -61 | -34 |
| Cerebellar-Basal Ganglia | 216 | -41 | -70 | -46 |
| Cerebellar-Basal Ganglia | 217 | -13 | 15 | 9 |
| Cerebellar-Basal Ganglia | 218 | -28 | 1 | 3 |
| Cerebellar-Basal Ganglia | 219 | -25 | 8 | 6 |
| Cerebellar-Basal Ganglia | 220 | -16 | -52 | -17 |
| Cerebellar-Basal Ganglia | 221 | -26 | -60 | -25 |
| Cerebellar-Basal Ganglia | 222 | -20 | 12 | -3 |
| Cerebellar-Basal Ganglia | 223 | -9 | 9 | 8 |
| Cerebellar-Basal Ganglia | 224 | -10 | 15 | 0 |
| Cerebellar-Basal Ganglia | 225 | 47 | -61 | -34 |
| Cerebellar-Basal Ganglia | 226 | 41 | -70 | -46 |
| Cerebellar-Basal Ganglia | 227 | 13 | 15 | 9 |
| Cerebellar-Basal Ganglia | 228 | 28 | 1 | 3 |
| Cerebellar-Basal Ganglia | 229 | 25 | 8 | 6 |
| Cerebellar-Basal Ganglia | 230 | 16 | -52 | -17 |
| Cerebellar-Basal Ganglia | 231 | 26 | -60 | -25 |
| Cerebellar-Basal Ganglia | 232 | 20 | 12 | -3 |
| Cerebellar-Basal Ganglia | 233 | 9 | 9 | 8 |
| Cerebellar-Basal Ganglia | 234 | 10 | 15 | 0 |

**Supplementary Table 1.** *MNI coordinates for ROIs used in network segregation*

*analyses.* Retrieved from Cassady et al. (2019) and Hausman et al. (2020).

| Effect | Dfn | Dfd | F | P | Effect Size |
| --- | --- | --- | --- | --- | --- |
| 2 (Reproductive vs Early Postmenopausal)  X 2 (Female vs Male) ANOVAs | | | | | |
| *All Networks* | | | | | |
| Stage | 1 | 185 | 1.19 | 0.278 | 0.01 |
| Sex | 1 | 185 | 0.28 | 0.598 | 0.00 |
| Stage:Sex | 1 | 185 | 0.05 | 0.818 | 0.00 |
| *Auditory* | | | | | |
| Stage | 1 | 186 | 0.04 | 0.836 | 0.00 |
| Sex | 1 | 186 | 4.96 | *0.027 | 0.03 |
| Stage:Sex | 1 | 186 | 0.31 | 0.576 | 0.00 |
| *Cerebellar-Basal Ganglia* | | | | | |
| Stage | 1 | 186 | 1.20 | 0.275 | 0.01 |
| Sex | 1 | 186 | 0.25 | 0.620 | 0.00 |
| Stage:Sex | 1 | 186 | 0.00 | 0.979 | 0.00 |
| *Cingulo-Opercular Task Control* | | | | | |
| Stage | 1 | 184 | 3.74 | 0.055 | 0.02 |
| Sex | 1 | 184 | 0.16 | 0.694 | 0.00 |
| Stage:Sex | 1 | 184 | 0.18 | 0.676 | 0.00 |
| *Default Mode* | | | | | |
| Stage | 1 | 186 | 6.01 | *0.015 | 0.03 |
| Sex | 1 | 186 | 0.86 | 0.355 | 0.00 |
| Stage:Sex | 1 | 186 | 0.55 | 0.461 | 0.00 |
| *Dorsal Attention* | | | | | |
| Stage | 1 | 186 | 4.16 | *0.043 | 0.02 |
| Sex | 1 | 186 | 0.13 | 0.724 | 0.00 |
| Stage:Sex | 1 | 186 | 5.69 | *0.018 | 0.03 |
| *Fronto-Parietal Task Control* | | | | | |
| Stage | 1 | 184 | 9.51 | **0.002 | 0.05 |
| Sex | 1 | 184 | 0.12 | 0.728 | 0.00 |
| Stage:Sex | 1 | 184 | 1.23 | 0.268 | 0.01 |
| *Salience* | | | | | |
| Stage | 1 | 185 | 2.69 | 0.102 | 0.01 |
| Sex | 1 | 185 | 0.05 | 0.823 | 0.00 |
| Stage:Sex | 1 | 185 | 0.02 | 0.885 | 0.00 |
| *Sensory Somatomotor Hand* | | | | | |
| Stage | 1 | 185 | 0.23 | 0.630 | 0.00 |
| Sex | 1 | 185 | 1.40 | 0.238 | 0.01 |
| Stage:Sex | 1 | 185 | 0.85 | 0.358 | 0.00 |
| *Sensory Somatomotor Mouth* | | | | | |
| Stage | 1 | 185 | 1.07 | 0.303 | 0.01 |
| Sex | 1 | 185 | 1.41 | 0.237 | 0.01 |
| Stage:Sex | 1 | 185 | 0.10 | 0.754 | 0.00 |
| *Ventral Attention* | | | | | |
| Stage | 1 | 185 | 0.88 | 0.349 | 0.00 |
| Sex | 1 | 185 | 4.29 | *0.040 | 0.02 |
| Stage:Sex | 1 | 185 | 4.02 | *0.047 | 0.02 |
| *Visual* | | | | | |
| Stage | 1 | 186 | 0.02 | 0.885 | 0.00 |
| Sex | 1 | 186 | 0.01 | 0.916 | 0.00 |
| Stage:Sex | 1 | 186 | 3.95 | *0.048 | 0.02 |

**Supplementary Table 2**. *Exploratory between-subjects ANOVA*

*results*. *: *p* < .05; **: *p* < .01; ***: *p* < .001.
